# Scraping the Surface: First Records of Cleaning Associations Between Sharks and Oceanic Manta Rays

**DOI:** 10.1101/2025.04.04.647128

**Authors:** Jane Vinesky, James Ketchum, Mauricio Hoyos

## Abstract

Interspecific scraping behaviors among large marine vertebrates may serve as an important mechanism for ectoparasite removal. Here, we present the first documented observations of shark–manta scraping interactions, which occurred in the Revillagigedo Archipelago, a remote marine protected area in the eastern tropical Pacific. Opportunistic video footage was collected at two dive sites between December 2024 and February 2025, capturing three discrete events in which Galapagos sharks (*Carcharhinus galapagensis*) initiated contact with three different oceanic manta rays (*Mobula birostris*). Using a focal-animal sampling approach, we quantified the number and rate of scraping events and categorized responses. Sharks scraped against mainly the ventral surfaces of mantas using their heads, gill regions, and lateral areas associated with high ectoparasite loads. Manta responses ranged from passive tolerance to active evasion, with the strongest reactions observed in response to the adult Galapagos shark. Scraping rates varied by site and shark size. These interactions occurred at or near established cleaning stations, suggesting that sharks may opportunistically use mantas as alternative cleaning substrates. While this may indicate behavioral plasticity among sharks, it raises concern about potential costs to mantas, particularly if some ectoparasites or pathogens are transferred. Our findings highlight the complexity of shifting ecological interactions among marine megafauna and underscore the importance of understanding behavioral responses to cleaning disruption for endangered oceanic mantas.

## Introduction

The Revillagigedo Archipelago is a remote group of four volcanic islands situated 386 km southwest of the Baja California Peninsula and approximately 800 km west of mainland Mexico (Aguirre-Muñoz et al., 2015) (Figure 1). Recognized as a UNESCO World Heritage Site in 2016 for its outstanding universal value to humanity and as a National Park in 2017, it provides an important refuge for migratory marine megafauna moving throughout the eastern tropical Pacific (UNESCO 2016; Lara-Lizardi et al., 2022).

**Figure 1.**
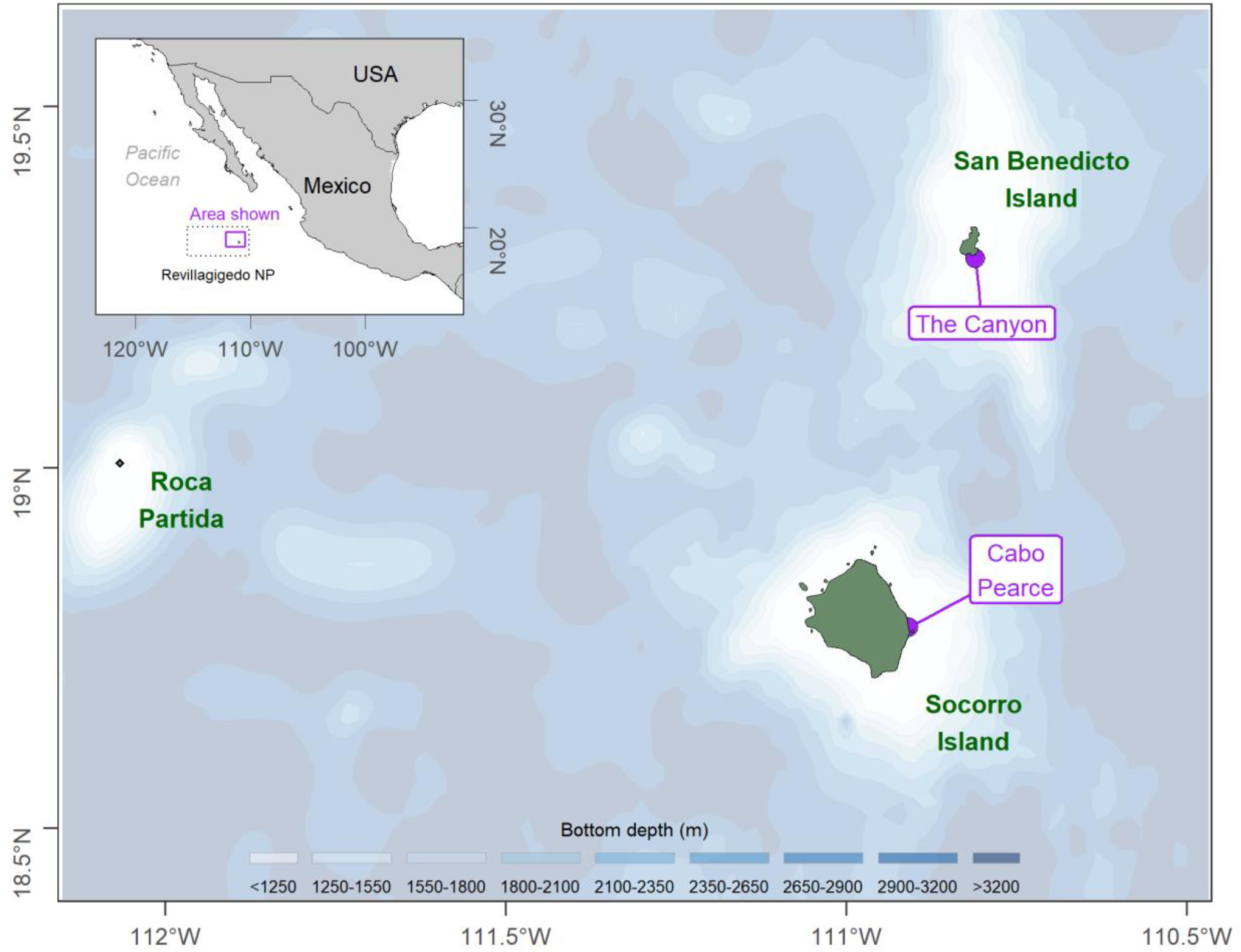
Locations where behavioral events were recorded (purple dots) within the Revillagigedo Archipelago.

Although remote, the archipelago’s fauna remains vulnerable to growing anthropogenic pressures, including illegal, unreported, and unregulated (IUU) fishing, plastic pollution, and aquarium trade (Clua et al., 2019; Pelamatti, 2021; Favoretto et al., 2023). The cumulative ecological impacts of these stressors remain poorly understood. Some ecological associations among marine megafauna and between elasmobranchs have been described within the Revillagigedo Archipelago (Becerril-García et al., 2019; Pancaldi, Becerril-García and Higuera-Rivas, 2022) along a multidimensional spectrum with overall fitness outcomes shifting between mutualism and parasitism (Gayford, 2024).

Oceanic manta rays (also known as mantas; *Mobula birostris*) are a flagship species in this ecosystem and are frequently observed aggregating at shallow cleaning stations throughout the archipelago, where they receive ectoparasite removal services from resident cleaner fish, notably the endemic Clarion angelfish (*Holacanthus clarionensis*) (Becerril-García et al., 2019; Bocos, AA, pers. comm.). These cleaning interactions are essential to manta health and may influence spatial use patterns and site fidelity (Murie, Spencer & Oliver, 2020). However, preliminary data suggest that populations of Clarion angelfish have declined in recent years (Clua et al. 2019, Bocos, AA, pers. comm.) potentially reducing the availability and effectiveness of cleaning services. At the same time, rising tourism pressure and diver presence at cleaning stations (Gómez-García et al. 2021) have been observed to disrupt cleaning events, raising concerns about cascading impacts on manta health and broader community dynamics.

In this context, interspecific scraping behavior between sharks and mantas presents a lens through which to examine behavioral flexibility in response to ecological disruption. While intraspecific and conspecific scraping has been documented among elasmobranchs (Kilmley et al. 2023), shark scraping on mantas remains unreported. These rare interactions provide a unique opportunity to explore the functional ecology of ectoparasite removal and the potential role of mantas as mobile substrates for other megafauna.

Here, we present the first documented observations of shark–manta scraping interactions, which occurred in the Revillagigedo Archipelago. We describe the context, frequency, and characteristics of these events, and assess the behavioral responses of mantas to shark-initiated contact. Our aim is to contribute to a broader understanding of how behavioral associations among marine megafauna arise in the context of anthropogenic stressors and environmental change.

## Methods

### 2.1 Study Area and Video Collection

This study was conducted at two dive sites in the Revillagigedo Archipelago, Mexico: The Canyon and Cabo Pearce (Figure 1). All observations were made during scuba dives between December 2024 and February 2025, with interactions recorded by divers using handheld GoPro cameras at depths between ∼4.5–20.5 meters. Video recordings were collected opportunistically, without the use of bait or any manipulation of shark or manta behavior (Gómez-García et al., 2021). Each video captured a discrete shark–manta interaction.

We employed an opportunistic focal-animal sampling approach (Altman, 1974), treating each video as a discrete behavioral observation centered on a single manta. The primary behavior of interest was shark scraping, defined as physical contact in which a shark rubs any part of its body against a manta. For each video, we recorded the shark species, body parts used to scrape, the location of contact on the manta (dorsal or ventral), the number of scraping events, and the manta’s behavioral response.

### 2.2 Behavioral Scoring and Analysis

Ethograms, or behavioral catalogs, are structured lists used in animal behavior studies to describe the most observed behaviors, classified as events or states, exhibited by animals under specific conditions (Altman, 1974; Gómez-García et al., 2021). Manta behaviors were categorized using an ethogram (Table 1), incorporating both functional (e.g., evasion, passive tolerance) and kinetic (e.g., acceleration, rolling) elements. To account for variability in video duration, we calculated scraping rates as events per minute categorized as discrete behaviors (Altman, 1974).

**Table 1.** Ethogram of behaviors observed during shark scraping interactions with mantas.

| Behavioral Unit | Definition | Type | Performer | Functional Category |
| --- | --- | --- | --- | --- |
| Head Scraping | Shark contacts manta with head (typically ventral area) | Event | Shark | Scraping |
| Lateral Body Scraping | Shark rubs flank or gill region against manta | Event | Shark | Scraping |
| Pre-contact Acceleration | Shark speeds up before initiating contact | Event | Shark | Pre-scraping |
| Post-contact Following | Shark follows closely after contact | State | Shark | Post-scraping |
| Passive Tolerance | Manta does not evade or reacts only with posture changes | State | Manta | Response (Tolerant) |
| Evasion | Manta accelerates or redirects to increase distance | State | Manta | Response (Avoidance) |
| Backward Roll | Manta performs tight rotation away from shark | Event | Manta | Response (Avoidance) |
| Cephalic Lobe Twitch | Small postural change or movement of cephalic lobes | Event | Manta | Response (Passive) |
| Return to Divers | Manta returns to observing divers after evading shark | Event | Manta | Post-contact |

Scraping was treated as a discrete behavior and its frequency (events per video) served as the primary metric. Behavioral states (e.g., evasion, directional swimming) were recorded descriptively and treated as time-linked responses rather than rate-based events.

## Results

We opportunistically documented three interactions between Galapagos sharks (*Carcharhinus galapagensis*) (n=3) and oceanic manta rays (*Mobula birostris*) (n=3) at two popular dive sites within the Revillagigedo Archipelago known as The Canyon and Cabo Pearce. All interactions occurred at or near known cleaning stations, and all were observed during recreational dives (∼4.5–20.5 m depth). In each case, the shark initiated the contact by scraping its head against a manta. Two interactions involved juvenile Galapagos sharks, and one involved an adult Galapagos shark. The total number of scraping events per interaction ranged from one to three, with manta responses varying by shark size (juvenile vs adult) and the number of scrape attempts.

Scraping was most frequently directed at the ventral surface of the mantas, particularly the abdominal region and anterior margin near the mouth. In all cases, the shark approached slowly and accelerated slightly prior to contact.

Mantas exhibited a range of behavioral responses (Table 1). In two of three interactions, the manta tolerated shark scraping with only minor postural adjustments such as cephalic lobe twitching or slight body angling, both classified as passive tolerance. In the remaining interaction, which occurred at The Canyon and involved an adult Galapagos shark, the manta exhibited strong evasive behavior following repeated scraping attempts including three backward rolls, multiple directional changes, and accelerations away from the shark, with the intensity of evasion increasing across the five scraping events. This was the only interaction in which evasion was observed following both the first and second scrapes. These findings suggest that shark size, persistence, or scrape frequency may influence manta responses. No mantas were observed to initiate contact, approach the shark, or return to the shark post-separation.

Across all interactions, Galapagos sharks scraped using their head, lateral sides, gill regions, and eye area (Figure 2, Panel B). Scraping rates ranged from 0.44 to 2.97 scrapes per minute, with the highest rate observed during the adult Galapagos shark interaction at The Canyon (2.97 scrapes·min^−1^). Lower rates were recorded in the two juvenile interactions with 0.58 scrapes·min^−1^at Cabo Pearce and 0.44 scrapes·min^−1^at The Canyon. These differences may reflect variation in individual motivation to scrape, opportunity for contact, or manta tolerance.

**Figure 2.**
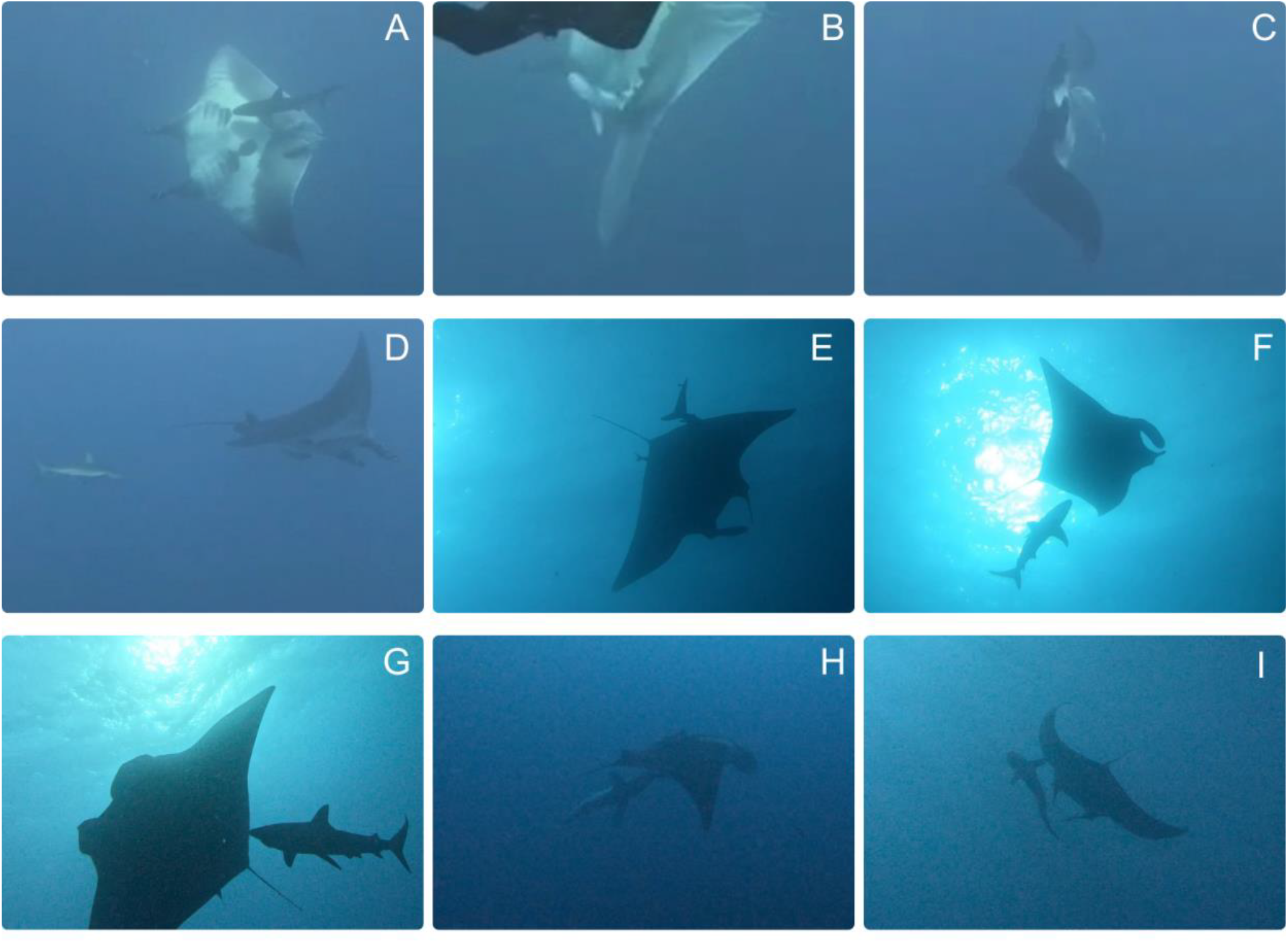
Representative images of behaviors portrayed in Table 1: A) Backward roll (evasion), B) Ventral scraping using dorsal head surface (shark scraping), C) Ventral scraping using lateral head surface (shark scraping), D) Post-contact following by shark; manta exhibits slight angular displacement (post-contact following, passive tolerance), E) Scrape initiation; manta swims upward (shark scraping, directional evasion), F) Close following prior to disengagement (post-contact following), G) Second scraping event; manta performs backward roll (shark scraping, evasion), H) Third contact followed by immediate directional change (shark scraping, evasion), I) Shark follows at short distance; manta swims forward (post-contact following, directional swimming).

## Discussion

This study provides novel observations of shark–manta interactions in the Revillagigedo Archipelago, documenting three discrete scraping events involving juvenile and adult Galapagos sharks. These interactions occurred in direct proximity to known cleaning stations, yet no cleaner fish were observed engaging with the mantas during the events, suggesting that sharks may opportunistically exploit mantas as alternative scraping substrates when traditional cleaning services are unavailable or disrupted.

Mantas exhibited variable responses to shark-initiated contact, ranging from passive tolerance to strong evasive maneuvers. Both interactions with juvenile Galapagos sharks elicited only mild or passive responses. In contrast, the single encounter with the adult Galapagos shark triggered a near immediate evasion followed by multiple backward rolls and withdrawal from the area. These behavioral differences may reflect species-specific risk perception by the mantas, particularly as some sharks have been documented preying on juvenile mantas (Strike et al., 2022), and the authors have observed shark bite scars on mantas in the region (Hoyos, MH, pers. obs.). Mantas can detect and respond to visual cues from potential threats at distances exceeding nine meters (Ari and Correia, 2008), and therefore the heightened response to the larger adult Galapagos shark may reflect a recognition of higher predation risk based on visual profiling or prior experience.

Variation in response may also be influenced by individual personality traits. Recent studies indicate that boldness and risk tolerance can vary among individuals, even in marine species, potentially affecting how animals respond to novel or potentially threatening stimuli (Wolf and Weissing, 2012). Though we could not identify individual mantas across videos, future studies employing photo-identification or tagging may provide insight into whether consistent behavioral types are present in these populations.

The differences in scraping rates across observations may be attributed to motivation to scrape, the availability of suitable body surfaces, or tolerance of the manta. Sharks scraped against mainly the ventral surfaces of mantas using their heads, gill regions, and lateral areas associated with high ectoparasite loads Unlike more stationary cleaning stations, where multiple clients may compete for attention, shark–manta interactions may represent a low-competition, mobile alternative when cleaner fish are absent or inactive. It is also possible that diver presence during the dives and safety stops may have influenced cleaner fish behavior, thereby increasing the likelihood of sharks seeking alternative mechanisms for ectoparasite removal.

Our findings add to a growing body of evidence that ecological associations among marine megafauna are dynamic, context dependent, and influenced by both environmental and individual level factors. In this case, mantas appear to function as an opportunistic substrate for scraping behavior, with the classification of the interaction shifting along a mutualism-parasitism spectrum depending on the species involved, the number of scrape attempts, and the behavioral response of the manta.

The unique behavioral ecology of mantas at Revillagigedo Archipelago may also contribute to the interaction dynamics observed. According to experienced divers and local environmental authority personnel, mantas at this site are highly acclimated to divers and frequently approach scuba bubbles, a behavior that could respond to a substitute-seeking response in the absence of traditional cleaning services. This interpretation is particularly relevant given recent anecdotal and survey-based evidence suggesting that populations of the primary cleaner symbiont of mantas in the region, the Clarion angelfish (*Holacanthus clarionensis*), have declined in recent years, potentially reducing cleaner availability (Clua et al., 2019, Bocos, AA, pers. comm.).

Concurrently, the Revillagigedo Archipelago has experienced a notable increase in marine tourism, with divers undertaking over 200 trips per season (November to June), often congregating at cleaning stations (E. Frias, pers. comm.). Despite briefing protocols and park regulations intended to minimize disturbance, cleaning behavior is frequently disrupted by divers (Gómez-García et al. 2021). This increasing anthropogenic pressure may be altering the functional ecology of cleaning stations, potentially driving both sharks and mantas to seek alternative cleaning strategies which may negatively affect them, such as tolerating contact from other megafauna like sharks, as compensatory behavior in a changing cleaning landscape.

While shark scraping may serve a functional role for the scraping species, the implications for mantas remain unclear. Galapagos sharks consistently scraped their heads, lateral flanks, gill regions, and eye areas, which are typically associated with high ectoparasite loads. These interactions could impose costs, particularly if sharks are transferring ectoparasites or microorganisms onto the mantas. Given the observed decline in Clarion angelfish and increasing disruption of traditional cleaning behavior, it is possible that mantas are becoming incidental hosts in a novel interaction that offers them little or no benefit and potentially increases their parasite burden and physiological stress. Future research should evaluate whether these interactions affect shark and manta health, either through pathogen transmission or behavioral stress, especially as cleaner fish communities decline in response to environmental and anthropogenic pressures.

## Acknowledgements

We would like to thank the citizen scientists whose footage made this work possible; Daniel Schuessler and Gabriele Hofstāter. We are grateful to Pacific Manta Research Group, The Manta Trust, Pelagios Kakunjá and Fins Attached for supporting us with expeditions. We also thank the Comisión Nacional de Áreas Naturales Protegidas (CONANP) and the Revillagigedo National Park for research permits and continued conservation efforts in the region.

## Notes

### Competing Interest Statement

The authors have declared no competing interest.

